# *Shigella sonnei* O-antigen inhibits internalisation, vacuole escape and inflammasome activation

**DOI:** 10.1101/799379

**Authors:** Jayne L Watson, Julia Sanchez-Garrido, Philippa J Goddard, Vincenzo Torraca, Serge Mostowy, Avinash R Shenoy, Abigail Clements

## Abstract

Two *Shigella* species, *flexneri* and *sonnei*, cause approximately 90% of bacterial dysentery worldwide. While *S. flexneri* is the dominant species in low-income countries, *S. sonnei* causes the majority of infections in middle and high-income countries. *S. flexneri* is a prototypic cytosolic bacterium; once intracellular it rapidly escapes the phagocytic vacuole and causes pyroptosis of macrophages, which is important for pathogenesis and bacterial spread. By contrast little is known about the invasion, vacuole escape and induction of pyroptosis during *S. sonnei* infection of macrophages. We demonstrate that *S. sonnei* causes substantially less pyroptosis in human primary monocyte-derived macrophages and THP1 cells. This is due to reduced bacterial uptake and lower relative vacuole escape, which results in fewer cytosolic *S. sonnei* and hence reduced activation of caspase-1 inflammasomes. Mechanistically, the O-antigen, which in *S. sonnei* is contained in both the lipopolysaccharide and the capsule, was responsible for reduced uptake and the T3SS was required for vacuole escape. Our findings suggest that *S. sonnei* has adapted to an extracellular lifestyle by incorporating additional O-antigen into its surface structures compared to other *Shigella* species.

## Introduction

*Shigella* are the causative agents of Shigellosis, infecting an estimated 125 million people annually. Children under five are most at risk with 61% of cases and 69% of deaths occurring in this age group (1). Closely related to *Escherichia coli*, the genus is made up of four species; *S. flexneri, S. sonnei, S. dysenteriae* and *S.boydii.* These are divided into serotypes based on the O-antigen (O-Ag) structure. *S. flexneri* and *S. sonnei* are responsible for the majority of infections however, dominance is highly dependent on the socioeconomic status of an area. *S. flexneri* is associated with poor water sanitation and hygiene in developing countries. In sub-Saharan Africa and Asia, *S. flexneri* accounts for 66% of cases and *S. sonnei* 24% of cases (2). However, in areas with good socioeconomic conditions and a high gross domestic product per capita, such as North America and Europe, *S. sonnei* is responsible for up to 80% of infections (3). Transitional countries that have recently undergone socioeconomic improvements show a shift from *S. flexneri* to *S. sonnei* as the dominant species (4-6). As a number of large populous countries undergo this shift (eg. Brazil, India, China), *S. sonnei* is emerging as an important pathogen.

The pathogenesis of *S. sonnei* is poorly understood and generally assumed to be similar to *S. flexneri*. The growing importance of *S. sonnei* has led to a re-evaluation of its pathogenesis and has revealed some important differences from *S. flexneri.* These include a novel adhesin, (7, 8), an antibacterial T6SS (9) and a group 4 capsule (G4C) which protects it from serum-mediated killing (10). Both species have a homologous T3SS that promotes secretion of effectors into host cells.

Unlike other *Shigella* species which contain multiple serotypes, there is only one *S. sonnei* serotype. The genes encoding biosynthesis and export of the O-Ag are encoded on the pSS virulence plasmid and were horizontally acquired from *Plesiomonas shigelloides*. In all other *Shigella* spp., these genes are located on the chromosome (11). The *S. sonnei* O-Ag is composed of two unusual sugars, 2-acetamido-2-deoxy-L-altruronic acid (L-AltNAcA) and 2-acetamido-2-deoxy-L-fucose (FucNAc4N), not present in the O-Ag of other *Shigella* spp., or indeed in many bacteria (12). Importantly, the G4C of *S. sonnei* is also composed of the O-Ag polysaccharide, linked to an unknown lipid anchor rather than the lipid A-core as in the LPS (10). Therefore the surface of *S. sonnei* is covered with two O-Ag layers.

Pyroptotic cell death is considered an important component of *S. flexneri* pathogenesis, (13) allowing *S. flexneri to* escape macrophage-mediated killing, induce local inflammation and invade epithelial cells from the basolateral side (14). In the canonical pathway for caspase-1 activation and pyroptosis, NOD and leucine-rich repeat containing proteins with CARD or PYD (NLRCs or NLRPs), AIM2-like receptors or Pyrin protein can respond to pathogen- and/or danger-associated molecular patterns. This leads to the assembly of the sensor e.g. NLRP3 or NLRC4, and the adaptor protein, ASC, into a signalling platform, known as the inflammasome, which activates caspase-1 (15). In the non-canonical pathway, caspase-4 directly senses and is activated by cytosolic LPS (16). Active-caspase-1 and active-caspase-4 can cleave gasdermin-D (GSDMD) (17). Once cleaved, the N-terminal of GSDMD forms pores in the cell membrane to cause swelling and membrane rupture. Proinflammatory cytokines, IL-1β and IL-18, are also cleaved by active caspase-1 into their mature forms and released (18, 19).

*S. flexneri* can activate the NLRC4 and NLRP3 inflammasomes (20). The T3SS needle and rod proteins (MxiH and MxiI, respectively) are recognised by hNaip/mNaip1 and mNaip2 proteins which interact with NLRC4 and promote caspase-1 activation (21, 22). NLRP3 senses decreased cytosolic potassium levels and activates caspase-1 (23). A T3SS effector, IpaH7.8, has been shown to be important for activation of both the NLRC4 and NLRP3 inflammasomes (20). In the case of *Shigella*, it is unclear whether pyroptosis benefits the host or the bacteria. *S. flexneri* is thought to use pyroptosis to escape the macrophage and infect epithelial cells. However, recent studies using *Salmonella* suggest that pyroptosis results in killing of bacteria by forming pore-induced intracellular traps (PITs) (24) or GSDMD targeting of bacterial membranes (25). It is currently unknown whether *S. sonnei* activates the same inflammasomes as *S. flexneri* and whether this is beneficial for the host or bacteria.

In this study, we demonstrate for the first time that *S. sonnei* induces caspase-1-dependent pyroptosis of human macrophages. However, we observed that equivalent bacterial inocula induced much less cell death for *S. sonnei* than *S. flexneri.* We show this is due to the O-Ag of *S. sonnei* which reduces internalisation and vacuole escape, resulting in less cytosolic bacteria. Our studies reveal an important role for the *S. sonnei* O-Ag in regulating bacterial interactions with macrophages, with one consequence being a reduction in inflammatory cell death.

## Results

### *S. sonnei* induces less macrophage cell death than *S. flexneri*

Previous research into the interactions of *Shigella* with macrophages has largely focused on *S. flexneri*, which robustly induces pyroptosis in macrophages (20). To investigate whether *S. sonnei* behaved in a similar manner, we infected primary human CD14+ monocyte derived macrophages (hMDMs) and measured the uptake of propidium iodide (PI), as an indicator of membrane damage that precedes pyroptosis. Unexpectedly, *S. sonnei* induced 50 % less PI uptake than *S. flexneri* (figure 1a).

**Figure 1.**
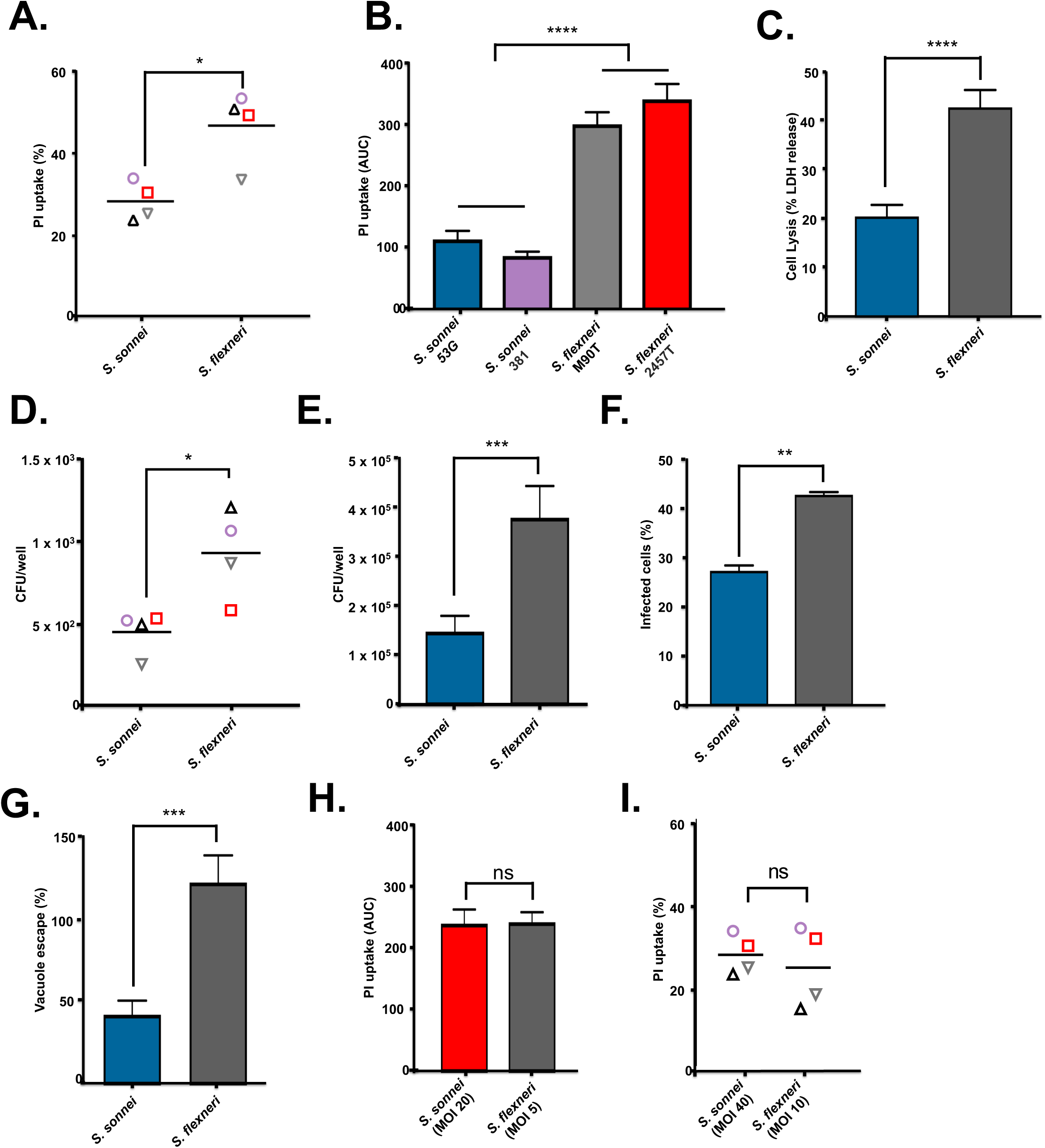
*S. sonnei* induces less pyroptosis of macrophages than *S. flexneri*. (A) Primary human monocyte-derived macrophage (hMDMs) were infected with the indicated *Shigella* strains and cell death was measured by PI uptake at 3 h post infection. *p<0.05 by paired Student’s t test, n=4 independent repeats from two donors. (B and C) THP1 cells were infected with the indicated wild type *Shigella* strains. Cell death was measured by PI uptake over a 3 h time-course and plotted as Area Under the Curve (AUC), n=3 (B) or LDH release at 3 h post infection, n=9 (C). ****p<0.0001 by one-way ANOVA with Tukey’s multiple comparisons test (B) or ****p<0.0001 by paired Student’s t test (C). (D) hMDMs and (E) THP1 cells were infected with *S. sonnei* or *S. flexneri* and gentamicin-protected intracellular bacteria determined by CFU enumeration. (D) *p<0.05 by paired Student’s t test, n=4 independent repeats from two donors and (E) ***p<0.001 by paired Student’s t test, n=11. (F) Immunofluorescence microscopy was used to visualise intracellular/extracellular bacteria and the percentage of infected THP1 cells calculated. **p<0.01 by paired Student’s t test, n=3. (G) THP1 cells were infected with *S. sonnei* or *S. flexneri* and subsequently treated with gentamicin alone or gentamicin and choloroquine to determine the percentage of cytosolic bacteria that have escaped the vacuole. ***p<0.001 by paired Student’s t test, n=4. (H) THP1 cells were infected with *S. sonnei* at an MOI of 20 or *S. flexneri* at an MOI of 5. Cell death was measured by PI uptake over a 3 h time-course and plotted as Area Under the Curve (AUC). ns, non-significant by paired Student’s t test, n=3. (I) hMDMs were infected with *S. sonnei* at an MOI of 40 or *S. flexneri* at an MOI of 10 and PI uptake was measured at 3 h post infection. ns, non-significant by paired Student’s t-test, n=4 independent repeats from two donors.

Similar experiments in PMA-differentiated THP1 cells recapitulated the reduced PI uptake during *S. sonnei* infection compared to *S. flexneri* (figure 1b), In addition, a lactate dehydrogenase (LDH) release assay comparing lytic cell death showed *S. sonnei* induced less cell death than *S. flexneri* (figure 1c). To ensure reduced cell death was not a unique feature of the widely used S. *sonnei* strain 53G, we included a recent clinical isolate, *S. sonnei* 381, alongside *S. sonnei* 53G and compared these to two different *S. flexneri* strains M90T (serotype 5a) and 2457T (serotype 2a). Notably, both *S. sonnei* strains induced lower PI uptake in macrophages (figure 1b).

### There are fewer cytosolic *S. sonnei* than *S. flexneri*

Induction of macrophage cell death by *S. flexneri* requires the bacteria to be cytosolic, which entails two steps: internalisation and vacuole escape. We hypothesised that differences in these processes between *S. flexneri* and *S. sonnei* might be responsible for the differences in cell death observed. To investigate why *S. sonnei* induced less cell death, we treated hMDMs or PMA-treated THP1 cells with 50 μM Z-VAD-fmk, a pan-caspase inhibitor, to inhibit cell death (figure S1a) and performed a gentamicin protection assay to calculate the number of intracellular bacteria (figure 1d and e). *S. sonnei* infected macrophages had reduced numbers of intracellular bacteria compared to *S. flexneri.*

As the earliest timepoint that can be measured in the gentamicin protection assay is 1 hour 40 min post infection, it is possible bacteria were already killed by this timepoint which would misrepresent the relative efficiency of internalisation (because internalised and killed bacteria would not be detected). To address this we enumerated intracellular bacteria by differential staining at 40 min post infection, which confirmed that fewer THP1 cells harboured intracellular bacteria when infected with *S. sonnei* than when infected with *S. flexneri* (figure 1f).

Internalised *S. flexneri* rapidly lyse the phagocytic/endosomal vacuole in order to access the cell cytosol and escape lysosomal degradation (26). To investigate how well *S. sonnei* escaped into the cytosol we used chloroquine (CHQ), an antibiotic that only accumulates in vacuoles at high enough concentrations to kill bacteria, allowing discrimination between cytosolic and vacuolar bacteria (27). *S. sonnei* showed a reduction in vacuole escape compared to *S. flexneri* (figure 1g). Taken together, these data indicated there are less cytosolic *S. sonnei* compared to *S. flexneri* at the same multiplicity of infection (MOI), which may result in the reduced macrophage cell death observed with *S. sonnei*. By increasing the *S. sonnei* MOI to obtain equivalent numbers of cytosolic bacteria to *S. flexneri* (figure S1b), *S. sonnei* and *S. flexneri* induced similar levels of cell death (figures 1h and j) and cell lysis (figure S1c). These findings confirm that cytosolic bacteria are required for induction of cell death in *S. sonnei* and *S. flexneri* and that *S. sonnei* does not access the cytosol as efficiently as *S. flexneri*.

### The T3SS is required for vacuole escape but not internalisation of *S. sonnei*

The T3SS of *S. flexneri* is required for bacteria to lyse the phagocytic vacuole and access the cytosol (28). Consistent with this, a *S. sonnei* T3SS mutant (Δ*mxiD*) had an impaired ability to escape the vacuole (figure 2a) and reduced cell death measured by PI uptake (figure 2b) and LDH release (figure S1d). The *S. sonnei* T3SS was required to induce vacuole lysis and hence produce cytosolic bacteria.

**Figure 2.**
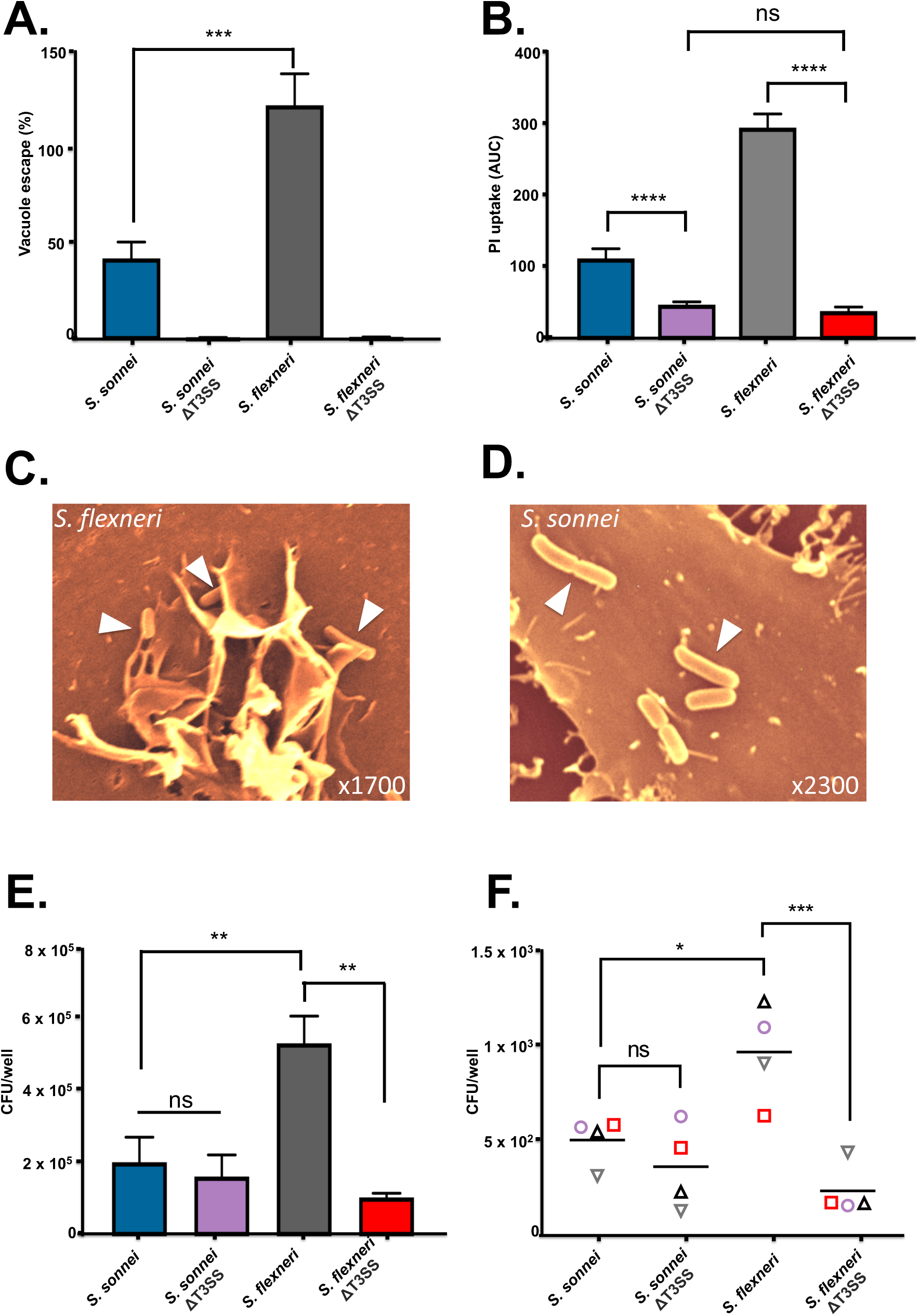
(A) THP1 cells were infected with *S. sonnei* or *S. flexneri* and their respective T3SS mutants and subsequently treated with gentamicin alone or gentamicin and choloroquine to determine the percentage of cytosolic bacteria that have escaped the vacuole. ***p<0.001 by one-way ANOVA with Tukey’s multiple comparisons test, n=4. (B) THP1 cells were infected with wild type *S. sonnei* and *S. flexneri* and their respective T3SS mutants. Cell death was measured by PI uptake over a 3 h time-course and plotted as Area Under the Curve (AUC). ns, non-significant, ****p<0.0001 by one-way ANOVA withTukey’s multiple comparisons test, n=3. (C and D) HeLa cells were infected with wild type *S. sonnei* and *S. flexneri* for 10 min before being washed and fixed for SEM analysis. Arrows indicate bacteria attached to the cell surface. (E) THP1 cells and (F) hMDMs were infected with wild type or T3SS-deficient *S. sonnei* and *S. flexneri* and gentamicin-protected internalised bacteria were determined by CFU enumeration. ns, non-significant, *p<0.05, **p<0.01 by one-way ANOVA with Tukey’s multiple comparisons test, (E) n=4 and (F) n=4 independent repeats from two donors.

It is unclear whether *Shigella* internalisation into macrophages is predominantly T3SS-dependent invasion or phagocytic uptake. T3SS-mediated invasion of epithelial cells by *S. flexneri* triggers extensive membrane recruitment to engulf the bacteria. To visualise *S. flexneri* and *S. sonnei* uptake we performed scanning electron microscopy (SEM) on infected cells and were able to see membrane recruitment around attached *S. flexneri* but not *S. sonnei* (figure 2c and d). As phagocytic uptake and T3SS-mediated invasion both involve membrane rearrangement these would be difficult to distinguish visually. Instead we performed gentamicin protection assays with wild type and T3SS mutants to quantify the number of intracellular bacteria in macrophages. Our experiments showed that in both hMDM and THP1 cells, internalisation into macrophages was T3SS-dependent for *S. flexneri* but not *S. sonnei* (figure 2e and f). This suggested that the majority of *S. flexneri* actively invaded macrophages, in contrast to *S. sonnei*, which were mainly internalised by phagocytic uptake.

### *S. sonnei* and *S. flexneri* induce similar pyroptosis pathways in infected macrophages

Given that cytosolic bacteria induce cell death through inflammasome activation we characterised the inflammasome pathways activated by *S. sonnei.* As the *S. flexneri* inflammasome activators MxiH, MxiI and IpaH7.8 proteins are 100 % identical between *S. sonnei* 53G and *S. flexneri* M90T we hypothesised they would activate the NLRC4 inflammasome. At comparable levels of cytosolic bacteria, similar activation of caspase-1 and proteolytic cleavage of GSDMD and IL-18 were observed (figure 3a). The involvement of the inflammasome pathway was confirmed using ASC^mRFP^ THP1 cells which revealed that both bacteria induced comparable levels of cells with ASC-containing inflammasome foci during infection (figure 3b and c). Further, infected GSDMD-silenced THP1 cells (THP^GSDMD-miR^, validated in S2a and b) underwent reduced cell death, suggesting pyroptosis is the dominant type of cell death induced by *S. sonnei* and *S. flexneri* (figure 3d).

**Figure 3.**
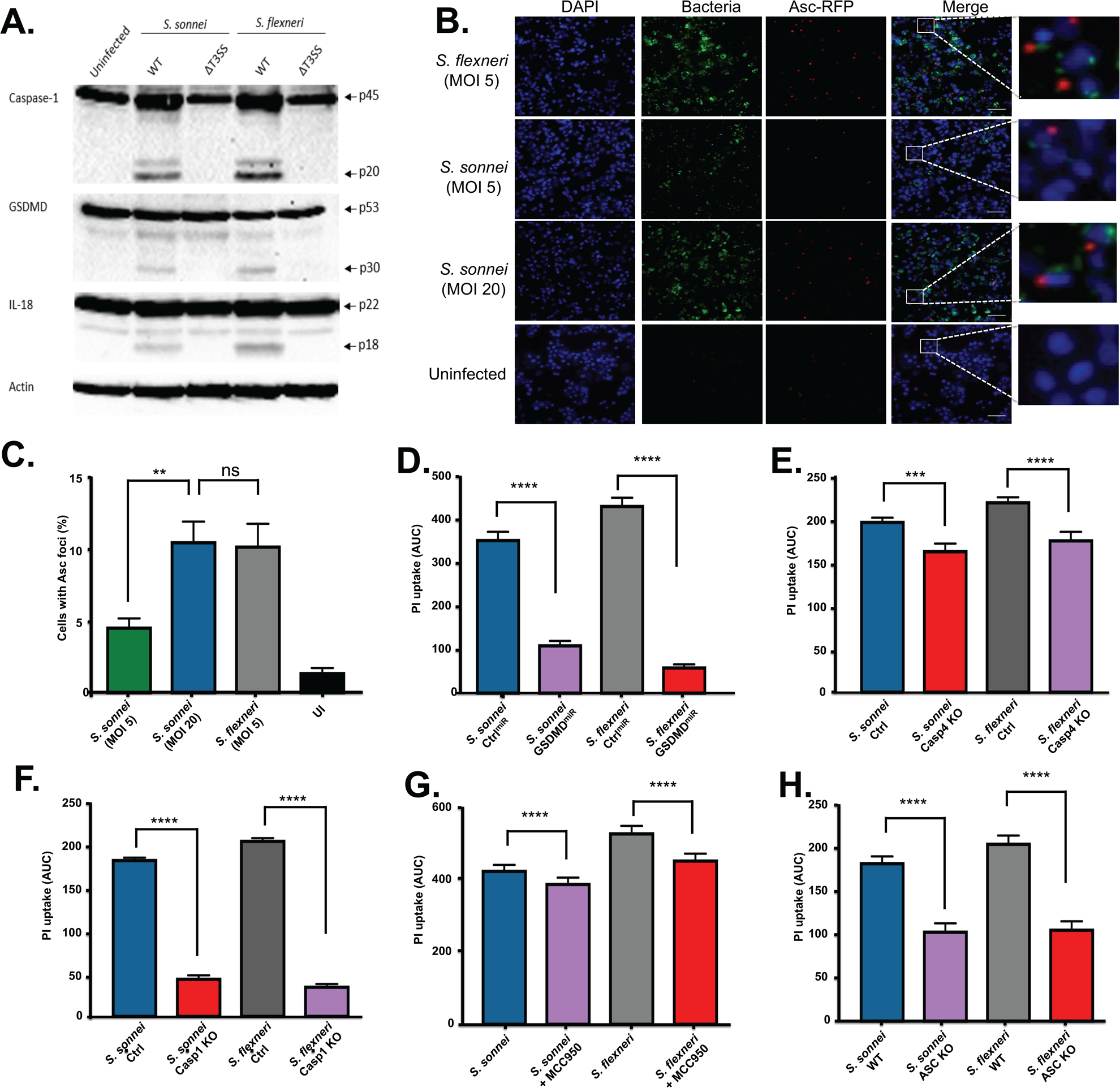
*S. sonnei* and *S. flexneri* induce similar pyroptotic pathways when normalised for numbers of cytosolic bacteria. (A) Immunoblots were performed on *Shigella-*infected THP1 cells to visualise cleavage of Caspase-1, GSDMD, and IL-18 at 3 h post infection. (B and C) ASC^mRFP^ THP1 cells were infected with fluorescent *Shigella* (green) at the indicated MOIs and after 3 h infection ASC foci formation visualised (red). DAPI staining was used to visualise DNA (blue). Representative micrographs for each strain are shown. The area indicated by the white box in the merged panel is enlarged to show single ASC foci in infected cells, and lack of ASC foci in uninfected cells. ASC foci formation was enumerated in (C). ns, non-significant, **p<0.01 by One-way ANOVA, n=4. (D) THP1 cells expressing non-targeting miRNA or GSDMD-targeting miRNA were infected with *S. sonnei (*MOI of 20) or *S. flex*neri (MOI of 5) for 3 h. Cell death was measured by PI uptake over a 3 h time-course and plotted as Area Under the Curve (AUC). ****p<0.0001 by one-way ANOVA with Tukey’s multiple comparisons test, n=3. (E) Control THP1 cells (Ctrl) and THP1 cells deficient for Caspase-4 (Casp4 KO) were infected with *S. sonnei* (MOI 20) or *S. flexneri* (MOI 5). Cell death was measured by PI uptake over a 3 h time-course and plotted as Area Under the Curve (AUC). ***p<0.001 and ****p<0.0001 by one-way ANOVA with Tukey’s multiple comparisons test, n=3. (F) Control THP1 cells (Ctrl) and THP1 cells deficient for Caspase-1 (Casp1 KO) were infected with *S. sonnei* (MOI 20) or *S. flexneri* (MOI 5). Cell death was measured by PI uptake over a 3 h time-course and plotted as Area Under the Curve (AUC). ****p<0.0001 by one-way ANOVA with Tukey’s multiple comparisons test, n=3. (G) THP1 cells left untreated or treated with 5 μM MCC950 were infected with *S. sonnei (*MOI of 20) or *S. flex*neri (MOI of 5) for 3 h. Cell death was measured by PI uptake over a 3 h time-course and plotted as Area Under the Curve (AUC). ****p<0.0001 by one-way ANOVA with Tukey’s multiple comparisons test, n=4. (H) THP1 cells and ASC-deficient THP1 cells were infected with *S. sonnei (*MOI of 20) or *S. flex*neri (MOI of 5) for 3 h. Cell death was measured by PI uptake over a 3 h time-course and plotted as Area Under the Curve (AUC). ****p<0.0001 by one-way ANOVA Tukey’s multiple comparisons test, n=3.

Cells deficient in caspase-4 showed reduced pyroptosis (figure 3e), however, loss of caspase-1 almost completely abolished pyroptosis (figure 3f, all knockout cells are validated in S2c -e), indicating the canonical pathway of pyroptosis predominates in *S. sonnei* and *S. flexneri* infected macrophages. Treatment with the NLRP3 inhibitor, MCC950 (29), did not markedly affect cell death (figure 3g and validated in S2f), suggesting that NLRP3 plays a minor role in pyroptosis. *ASC*-deficient THP1 cells showed a partial reduction in cell death levels compared to WT THP1 cells (figure 3h). Taken together, these results are consistent with NLRC4 activation contributing to pyroptosis during *S. sonnei* infection of human macrophages, which is similar to previous reports for *S. flexneri*.

### The T6SS and LVP instability do not account for reduced cell death caused by *S. sonnei*

All *Shigella* spp. harbour a large virulence plasmid (LVP) that encodes the T3SS, its effectors and additional important virulence factors. The LVP of *S. sonnei* is less stable than *S. flexneri* due to the evolution of different toxin:anti-toxin systems (30). We inserted an antibiotic resistance cassette onto the LVP to create a stabilised LVP and used this strain to test if LVP loss was affecting the amount of cell death that was induced. The LVP stabilised *S. sonnei* induced similar cell lysis as WT *S. sonnei* indicating that differences in plasmid retention was not responsible for the altered interaction with macrophages (figure 4a).

**Figure 4.**
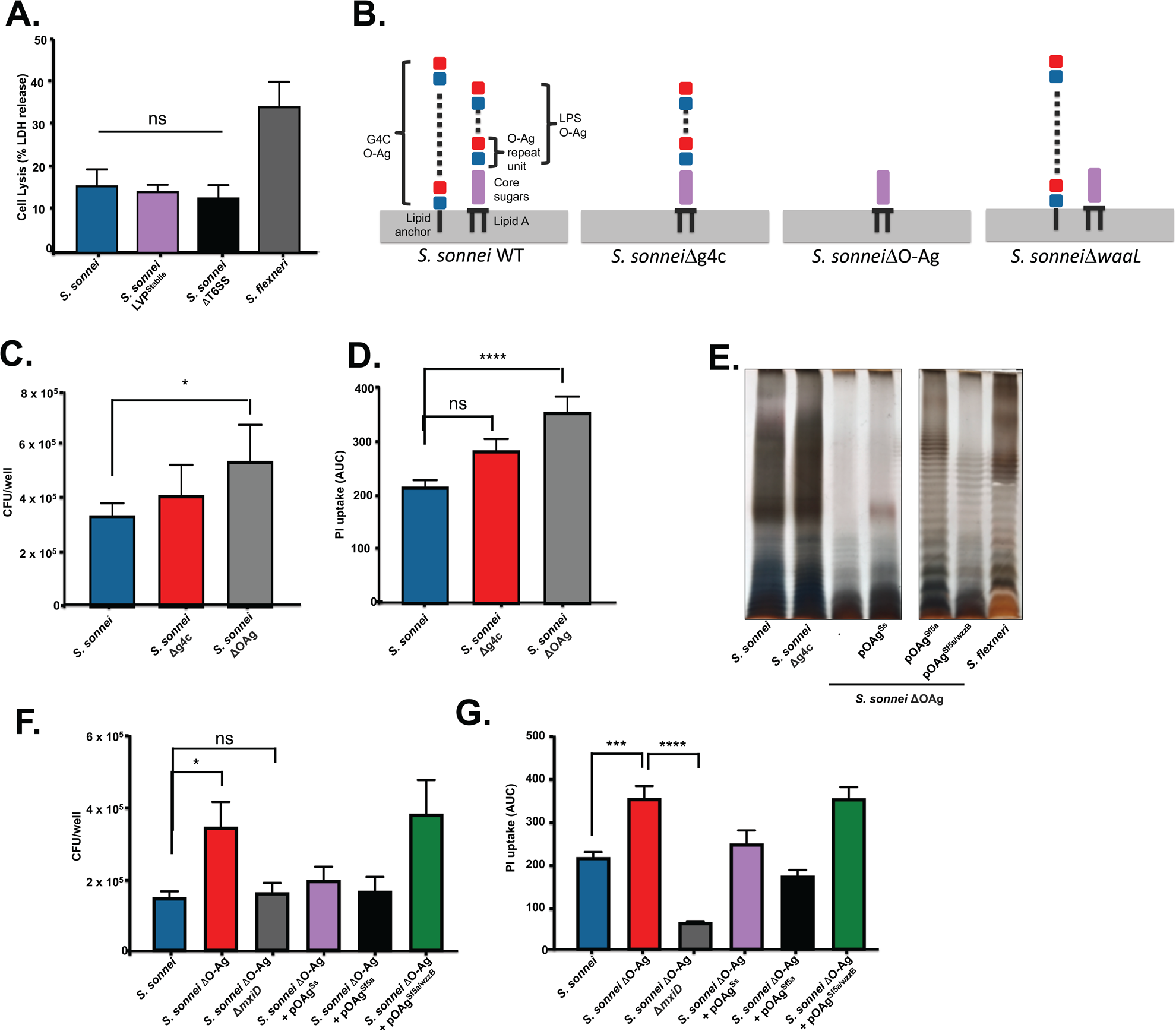
Presence of the *S. sonnei* O-Ag reduces cells death and internalisation. (A) THP1 cells were infected with wild type *S. sonnei, S. sonnei* LVP^Stabile^ or *S. sonnei* ΔT6SS and cell death was measured by LDH release at 3 h post infection. Ns, non-significant by one-way ANOVA with Tukey’s multiple comparisons test, n=3. (B) *S. sonnei* has O-Ag attached to the cell surface in two different forms, as the conventional O-Ag attached to the Lipid A-core of the LPS, and as an O-Ag capsule attached to the cell surface by an unknown lipid anchor. *S. sonnei* ΔG4C has only the O-Ag incorporated into the LPS, *S. sonnei* ΔO-Ag has neither G4C or LPS O-Ag and *S. sonnei ΔwaaL* has only the G4C O-Ag. (C) THP1 cells were infected with wild type *S. sonnei, S. sonnei* ΔG4C or *S. sonnei* ΔO-Ag and gentamicin-protected internalised bacteria were determined by CFU enumeration. *p<0.05 by one-way ANOVA with a mixed effects model with Sidak’s multiple comparisons test, n=10 (wild type) and 7 (mutants). (D) THP1 cells were infected with wild type *S. sonnei, S. sonnei* ΔG4C or *S. sonnei* ΔO-A*g*. Cell death was measured by PI uptake over a 3 h time-course and plotted as Area Under the Curve (AUC). ****p<0.0001 by one-way ANOVA with Tukey’s multiple comparisons test, n=4. (E) Crude LPS was purified from the indicated *S. sonnei* and *S. flexneri* strains, separated by 12% SDS-PAGE and visualised by modified silver stain. (F) THP1 cells were infected with wild type *S. sonnei, S. sonnei* ΔO-Ag or *S. sonnei* ΔO-Ag complemented with the *S. sonnei* O-Ag (pO-Ag^Ss^), the *S. flexneri* O-Ag (pO-Ag^Sf5a^) or the *S. flexneri* O-Ag and WzzB (pO-Ag^Sf5a/wzzB^) and gentamicin-protected internalised bacteria were determined by CFU enumeration. *p<0.05, **p<0.01 by one-way ANOVA with Tukey’s multiple comparisons test, n=4. (G) THP1 cells were infected with wild type *S. sonnei, S. sonnei* ΔO-Ag, *S. sonnei* ΔO-Ag + pO-Ag^Ss^ or *S. sonnei* ΔO-Ag + pO-Ag^Sf5a^. Cell death was measured by PI uptake over a 3 h time-course and plotted as AUC. ****p<0.0001 by one-way ANOVA with Tukey’s multiple comparisons test, n=4.

Even though the T6SS of *S. sonnei* has only been described to have anti-bacterial activity (9), T6SSs from other bacteria (e.g. *Francisella tularensis* (31, 32)) have activity within macrophages. We therefore created a *S. sonnei* T6SS mutant (Δ*tssB*) to determine if there was any contribution by the T6SS to cell death but found no difference in LDH release (figure 4a), indicating that the T6SS was not responsible for the altered interaction with macrophages. Altogether, these results ruled out loss of LVP or a contribution by the T6SS in the reduced cell death observed for *S. sonnei.*

### The *S. sonnei* O-Antigen prevents internalisation into macrophages

In *S. sonnei* the O-Ag is incorporated into the G4C as well as being attached to the lipid A-core of LPS (figure 4b). The incorporation of the O-Ag into LPS and G4C is genetically separable, which we exploited to investigate their respective roles in the interaction with macrophages. The G4C of *S. sonnei* reduces bacterial invasion of epithelial cells by impairing T3SS activity (10), and could therefore play a similar role in macrophage internalisation. We confirmed that *S. sonnei* ΔG4C invaded HeLa cells more efficiently (figure S3b). Uptake and pyroptosis induced by *S. sonnei* ΔG4C was statistically similar to wildtype bacteria, although we did observer slightly higher cell death with the Δg4c mutant (figure 4c-d). This was consistent with predominantly phagocytic uptake of *S. sonnei* by macrophages.

We then deleted the O-Ag synthesis operon (genes *wbgT* to *wbgZ*) (33) to create a strain devoid of all O-Ag (both LPS and G4C linked) (figure 4b). This strain (ΔO-Ag) demonstrated increased internalisation and cell death as compared to *S. sonnei* Δg4c (figure 4c and d) and wild type *S. sonnei*. By contrast, an LPS O-Ag deficient strain (Δ*waaL*), which retains the G4C, showed equivalent internalisation as wild type *S. sonnei* (figure S3a). Therefore, the presence of the *S. sonnei* O-Ag per se, rather than specifically the O-Ag in the capsule or LPS, impedes macrophage internalisation and its complete removal enhances bacterial internalisation.

We have shown that *S. sonnei* cell death is T3SS-dependent due to the requirement for cytosolic bacteria. The T3SS tip accessibility has previously been shown to be enhanced upon removal of the G4C, and further exposed by removal of the O-Ag (10). We therefore hypothesised that the O-Ag was impeding T3SS mediated invasion. To test this we created a T3SS mutant in the O-Ag deficient strain (ΔO-AgΔ*mxiD*). In keeping with our hypothesis this strain had wild-type levels of internalisation (figure 4e), but impaired activation of cell death because it is unable to escape the vacuole. To further investigate the role of the O-Ag in shielding the T3SS we complemented the O-Ag mutant with either the *S. sonnei* O-Ag synthesis operon (pSS) or the *S. flexneri* 5a O-Ag synthesis and modification operons (pSf5a) (34-36) (figure 4e). Both complemented strains impeded internalisation of *S. sonnei* (figure 4f) and, as a consequence, reduced the level of cell death similar to those observed with wildtype *S. sonnei* (figure 4f). Interestingly, complementation with pSf5a produced a *S. flexneri*-like O-Ag ladder which migrated differently on SDS-PAGE than when expressed in *S. flexneri*. To determine if this was due to different modal length of O-Ag controlled by WzzB we introduced the *wzzB*^Sf^ onto the pSf5a complementation plasmid (pSf5a/*wzzB*). In this strain the modal length of the O-Ag resembled that of the wild type *S. flexneri*, however the levels of internalisation and cell death were not reduced to the levels of the wild type *S. sonnei* and instead resembled the levels of the O-Ag mutant.

## Discussion

*S. flexneri* is known to induce pyroptosis in macrophages. This is considered a key step in the pathogenesis of *Shigella* as it allows bacteria to infect epithelial cells from the preferred basolateral side and leads to bacterial dissemination. In addition, pyroptosis creates an inflammatory response causing the recruitment of neutrophils, which disrupt the epithelial cell barrier and allows more *Shigella* to traverse the epithelial layer (37).

Here we present evidence that *S. sonnei* does not use the same mechanisms during infection as *S. flexneri* (summarised in figure 5). In line with previous reports, we found that *S. flexneri* induces rapid pyroptosis upon internalisation of infected macrophages (20, 22). However, *S. sonnei* induced markedly less macrophage cell death, which was the result of a decreased number of cytosolic bacteria through a combination of fewer internalised *S. sonnei* and impaired vacuole escape. The requirement for cytosolic bacteria in the induction of inflammasomes was consistent for both *S. sonnei* and *S. flexneri*. Additional host responses are also likely to be affected by the reduced number of cytosolic bacteria for *S. sonnei* compared to *S. flexneri*.

**Figure 5.**
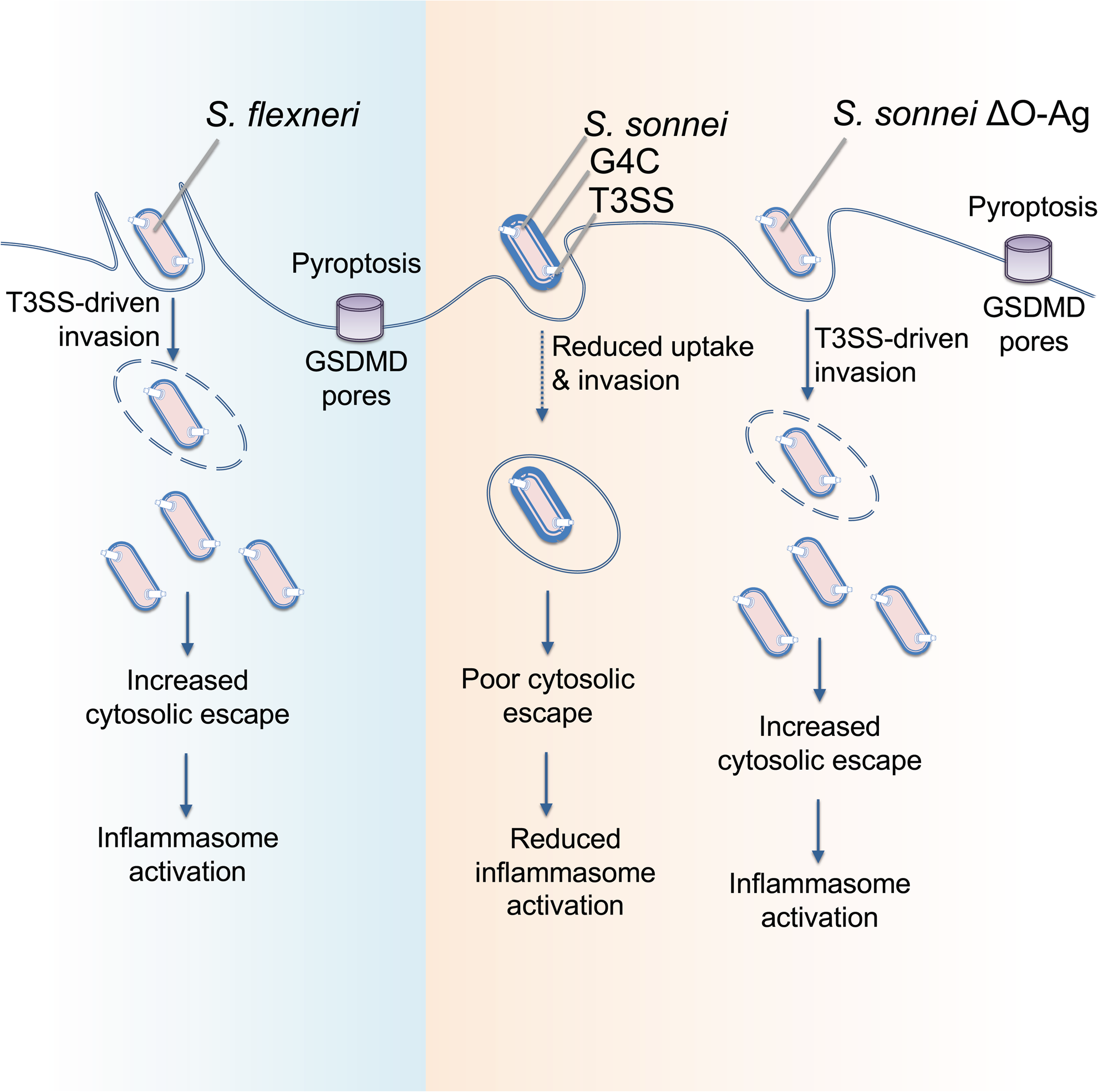
Model showing the interaction of *S. flexneri, S. sonnei* and *S. sonnei* ΔO-Ag with macrophages. *S. flexneri* uses T3SS-mediated invasion and vacuole rupture to reach the cytosol where it induces inflammasome activation, GSDMD pores and pyroptosis. *S. sonnei* does not invade macrophages using its T3SS and only becomes intracellular through phagocytosis. Combined with a decreased ability to escape the vacuole this reduces the number of cytosolic bacteria and leads to reduced inflammasome activation. The T3SS becomes accessible in *S. sonnei* ΔO-Ag allowing the bacteria to invade macrophages and reach the cytosol where they can activate the inflammasome and cause pyroptosis.

Once *S. sonnei* and *S. flexneri* cytosolic numbers were normalised, pyroptosis proceeded via similar pathways and to similar levels. For both species, cell death was predominantly dependent on GSDMD and caspase-1, indicating the canonical inflammasome pathway is induced by *Shigella.* There may be a minor contribution to cell death for the non-canonical pathway as immunoblots indicated that caspase-4 was activated by infection of both *S. sonnei* and *S. flexneri*, and caspase-4 deficiency or NLRP3 inhibition led to less pyroptosis over time than control cells. However, this difference was minor compared to that observed for ASC or caspase-1-deficient cells. NLRC4 has a caspase-recruitment and activation domain (CARD) which can enable its interaction with, and activation of caspase-1 directly, bypassing the need for ASC (38-40). This suggests that the NLRC4 inflammasome has a prominent role in the cell death of *S. sonnei*-infected THP1 cells. These results are in line with those shown previously for *S. flexneri*, which suggest both the NLRP3 and NLRC4 inflammasomes are involved in *S. flexneri*-mediated macrophage death (20).

Interestingly, *S. sonnei* was able to reduce internalisation into macrophages in an O-Ag dependent manner. The O-Ag contributes to host immune evasion and its role in evasion of complement mediated killing is well characterised (41). There are also examples of O-Ags affecting cellular interactions including impeding recognition and internalisation by epithelial cells (*Salmonella* Typhimurium (42)) and macrophages (*Burkholderia cenocepacia* (43)). The modal length of the O-Ag from *Salmonella* Typhimurium or *S. flexneri* serotype 2a is important for T3SS-mediated invasion into macrophages and epithelial cells respectively (44). Similarly, glucosylation of the *S. flexneri* serotype 5a O-Ag which reduces the O-Ag length by half enhances its invasiveness (45).

Unexpectedly, in our study the internalisation of *S. sonnei* into macrophages was independent of its T3SS. This is in contrast to *S. flexneri* which exhibits significant T3SS-mediated invasion into macrophages. This suggests that macrophage internalisation is a combination of bacteria-driven invasion and phagocytic uptake for *S. flexneri*, while *S. sonnei* internalisation is almost exclusively due to phagocytic uptake. The *S. sonnei* O-Ag is incorporated into both the G4C and the LPS of *S. sonnei*. Only when all the O-Ag layers of *S. sonnei* are removed is *S. sonnei* able to efficiently invade macrophages. The accessibility of IpaB was previously shown to increase upon removal of the G4C and a further increase was observed for an O-Ag-deficient strain indeed suggesting the lipid A-core linked O-Ag also contributes to shielding of the T3SS (10). The ability of the serotype 5a O-Ag synthesis and modification operon from *S. flexneri* to prevent internalisation of the O-Ag deficient *S. sonnei* indicates the composition of the saccharides are not important for this phenotype. Furthermore, the inability of the *S. flexneri* O-Ag when regulated by *wzzB* to complement for internalisation of cell death suggests the modal length of the O-Ag is important. However, this strain also produced a low amount of O-Ag and we cannot discount this as the reason for the failure to complement. Our data, and previously published data regarding the accessibility of the T3SS, supports the conclusion that the O-Ag acts as a physical barrier to T3SS-mediated invasion rather than being anti-phagocytic.

The results presented here, combined with previous investigations, indicate that *S. sonnei* and *S. flexneri* use different infection mechanisms. These mechanisms are also different from related Gram-negative enteric pathogens such as *Salmonella* or EPEC, which also activate distinct inflammasome pathways in human macrophages (46-49). Increasing evidence points to *S. sonnei* being more adapted to an extracellular lifestyle as, compared to *S. flexneri*, it invades epithelial cells and macrophages poorly. This may partly explain the dominance of *S. sonnei* in developed countries where improved living conditions, including reduced overcrowding and hence person-person spread of pathogens, fails to lower *S. sonnei* infection rates. These studies highlight that further investigation into *S. sonnei* is required in order to implement appropriate measures to reduce infection rates.

## Materials and Methods

### Bacterial strains and growth

Unless otherwise stated, all *Shigella* strains (Table S1) were routinely grown in tryptone soya broth (TSB) at 37°C with shaking at 200 RPM. Antibiotic selection was used when necessary as follows: 100 µg/mL ampicillin (Amp), 50 µg/mL kanamycin (Kn), 12.5 µg/mL chloramphenicol (Cm), 100 µg/mL erythromycin, 50 µg/mL streptomycin (Sm), 10 µg/mL gentamicin (Gm).

### Cloning and mutagenesis

*S. sonnei* LVP^Stabile^, *S. sonnei* Δ*waaL, S. sonnei* Δ*tssB* and *S. sonnei* ΔO-Ag strains were constructed as follows, primer sequences are in Table S2. For *S. sonnei* LVP^Stabile^ nt 82936-83715 and nt 83716-84215 were amplified using primers 1 and 2, and 3 and 4. The chloramphenicol cassette was amplified from pKD3 using primers 21 and 22. Overlapping PCR was used to construct the mutagenesis fragment consisting of 82936-83715-Cm-83716-84215, (note the P1-P2 fragment was inserted in the opposite orientation). This fragment was further amplified by PCR with primers 1 and 4. 2 μg of PCR product was electroporated into *S. sonnei* 53G + pKD46 induced with 1mM L-arabinose for 45 min to express lambda red recombinase genes. The electroporation was plated on TSB supplemented with Cm. Genomic insertion of *cat* was verified by PCR using primers 5 and 22

For *S. sonnei* Δ*tssB* 500bp fragments flanking *tssB* were amplified using primers 6 and 7, and 8 and 9. The kanamycin cassette was amplified from pKD4 using primers 21 and 22. Overlapping PCR was used to construct the mutagenesis fragment consisting of 5’ *tssB*-kan-3’ *tssB*. This fragment was further amplified by PCR with primers 6 and 9. 2ug of PCR product was electroporated into *S. sonnei* 53G + pKD46 induced with 1mM L-arabinose for 45 min to express lambda red recombinase genes. The electroporation was plated on TSB supplemented with Kn. Genomic insertion of *kan* was verified by PCR using primers 10 and 22.

For *S. sonnei* Δ*waaL* 500bp fragments flanking *waaL* were amplified using primers 11 and 12, and 13 and 14. The kanamycin cassette was amplified from pKD4 using primers 21 and 22. Overlapping PCR was used to construct the mutagenesis fragment consisting of 5’ *waaL*-kan-3’ *waaL*. This construct and pSEVA612S were digested with BamHI and EcoRI, ligated and transformed into *E. coli* CC118-λpir. The resulting plasmid pSEVAΔwaaL-Kn was conjugated into *S. sonnei* 53G. Briefly, 20 μl helper *E. coli* 1047 pRK2013 was incubated for 2h at 37°C with 20 μl of the donor strain (*E. coli* CC118-λpir pSEVAΔwaaL) on LB agar. 40 μl of the receiver strain (*S. sonnei* 53G with pACBSR) was added and the plate incubated for 4 h at 37°C. Conjugants were selected on TSB agar supplemented with Gm and Sm. Individual colonies were grown in TSB supplemented with Sm and 0.4% (w/v) L-arabinose (Sigma) for 8 h to induce expression of the I-SceI endonuclease from pACBSR, and plated on Kn plates. Genomic deletion of *waaL* was verified by PCR using primers 15 and 22. The strains were passaged several times in liquid TSB to remove pACBSR and bacteria sensitive to Sm were selected.

*S. sonnei* ΔO-Ag mutant was constructed by amplifying 500bp fragments upstream of *wbgT* and downstream of *wbgZ* using primers 16 and 17 and 18 and 19. The kanamycin cassette was amplified from pKD4 using primers 21 and 22. Overlapping PCR was used to construct the mutagenesis fragment consisting of 5’ *wbgT* -kan-3’ *wbgZ*. This construct and pSEVA612S were digested with BamHI and EcoRI, ligated and transformed into *E. coli* CC118-λpir. The resulting plasmid pSEVAΔO-Ag-Kn was conjugated into *S. sonnei* 53G as described above. Genomic deletion of ΔO-Ag was verified by PCR using primers 20 and 22.

Complementation vectors were constructed using standard molecular biology techniques. The 53g O-Ag operon was amplified with primers 23 and 24. The PCR product and pSEVA471 were digested with BamHI and ligated to create pO-Ag^Ss^. The M90T gtr operon was amplified with primers 25 and 26. The PCR product and pSEVA471 were digested with KpnI and BamHI and ligated to create pSEVA471-gtr. The M90T O-Ag operon was amplified with primers 27 and 28. The PCR product and pSEVA471-gtr were digested with BamHI and XbaI and ligated to create pOAg^Sf5a^. The M90T wzzB gene was amplified with primers 29 and 30. The PCR product and pOAg^Sf5a^ were digested with KpnI and ligated to create pOAg^Sf5a/*wzzB*^. All complementation constructs include predicted promoters and terminators.

### Cell culture and infection

THP-1 cells were maintained in Roswell Park Memorial Institute (RPMI) medium supplemented with 10% heat-inactivated foetal bovine serum (FBS), 5 mM Hepes, 5 mM sodium pyruvate, 100 μg/ml penicillin and 100 μg/ml streptomycin. Cells were seeded at 7.5×10^5^ cells/ml 72 hours prior to infection in complete RPMI + 100 ng/ml phorbol-12-myristate-13-acetate (PMA). 24 hours prior to infection, media was replaced with phenol-red free, PMA-free complete RPMI medium. HeLa cells were maintained in Dulbecco’s Modified Eagle’s Medium (DMEM) (1000 mg/L glucose) supplemented with 10 % FBS. Cells were seeded at 1×10^5^ cells/ml, 24 hours prior to infection. All cell lines were incubated at 37°C, 5% CO2. Cells were infected with indicated multiplicity of infection (MOI) and centrifuged for 10 mins at 600 xg to synchronise infection. At 30 minutes post centrifugation, Gm (150 μg/ml) was added to directly to wells for the remainder of the experiment. Where indicated, inhibitors, Z-VAD-fmk (50 μM; R&D) or MCC950 (5 μM; Tocris Bioscience), were added to cells 1 hour prior to infection. To induce NLRP3-driven caspase-1 activation, cells were primed with ultrapure O111:B4 LPS (250 ng/mL; Invivogen) for 3 hours and then treated with nigericin (20 μM; Sigma) for 45 minutes. To induce caspase-4 activation, unprimed cells were transfected with LPS (5 μg/ml) using Lipofectamine 2000 (1% v/w; Invitrogen).

HeLa infected cells were washed and fixed in 2.5% glutaraldehyde for analysis by scanning electron microscopy at an accelerating voltage of 25kV using a JEOL JSM-5300 scanning electron microscope [JEOL (UK), Herts, UK].

### Generation of cell lines

The THP1 GSDMD^miR^ cell line was generated previously as described (50), THP1 Casp1 KO, THP1 Casp4 KO, and THP1 ASC KO were all kindly provided by Veit Hornung (51).

### Isolation of primary human monocyte derived macrophages

Leukocytes cones were obtained from the NHS blood and transfusion service (from anonymous healthy donations), as previously described (46). Blood from each donor was diluted 1:4 with PBS, transferred into a LeucoSep tube (Greiner Bio-One) and centrifuged at 1000 xg, 20 mins at RT (slow acceleration and deceleration to prevent disturbance of the layers) to obtain the buffy coat containing white blood cells. This was separated and washed three times with RPMI. Cell were washed with MACS buffer (50 mg/ml BSA, 2 mM EDTA in PBS). CD14+ cells were isolated by MACS using biotinylated anti-CD14+ antibody and anti-biotin microbeads according to the manufacture’s protocol (Miltenyi Biotec). Monocytes were cultured in complete RPMI plus 20 ng/ml recombinant human M-CSF for 7 days to promote differentiation into hMDMs. Media was replaced with complete RPMI lacking antibiotics and M-CSF 24 hours prior to infection.

### Internalisation and vacuole-escape assays

To prevent cell death, cells were treated with Z-VAD-fmk (50 μM) 1 hour prior to infection. Cells were infected with bacteria as described above. For internalisation assays, cells were washed with serum-free RPMI and lysed with TritonX-100 (0.5%) at 1.5 hours post infection. For vacuole-escape assays, cells were treated 30 min post-infection with either 200 μg/ml chloroquine and 150 μg/ml gentamicin or 150 μg/ml gentamicin alone for 1 h and then lysed with TritonX-100 (0.5%). Serial dilutions were performed, plated on LB agar and incubated overnight at 37°C.

### PI uptake assays

Cells and bacterial strains were prepared as described above. Prior to infection, cells were supplemented with 5 μg/ml Propidium iodide (PI; Invitrogen). For timecourse assays, fluorescence was measured at 630 nm every 10 mins with POLARStar 623 Omega plate reader (BMG Labtech) (52). Uninfected controls treated with TritonX-100 (0.05%) were used to calculate percentage uptake.

### LDH assays

Infections were performed as described above. At 3 hours post infection, supernatants were harvested. An LDH assay was performed as per kit instructions (CytoTox 96® Non-Radioactive Cytotoxicity Assay, Promega). Absorbance was measure at 490 nm and values are expressed as percentage of 100% lysis control. All values are normalised to uninfected control.

### Immunoblots

Infections were performed as described previously, except prior to infection cells were washed with PBS and infections were done in OptiMEM + 5 mM Na pyruvate. Supernatants were precipitated in acetone (1:4 v/v) overnight at −20°C, acetone was aspirated, and samples left to air dry. Cells were lysed in RIPA buffer (120 mM Tris pH 8.0, 300 mM NaCl, 2% NP-40, 1% Na deoxycholate, 2 mM EDTA) supplemented with complete protease inhibitor and 1 mM PMSF. Laemmli buffer and 5% 2-mercaptoethanol were added to lysates. Precipitated supernatants were re-suspended in respective cell lysates to create pooled samples. Mouse anti-hcaspase-1 (AdipoGen), mouse anti-caspase-4 (Santa Cruz biotechnology), goat anti-hIL1β (R and D systems), rabbit anti-hIL18 (MBL international) were used at 1:1000 dilution and mouse anti-hGSDMD (Santa Cruz Biotechnology) was used at 1:500.

### Immunofluorescence microscopy

Cells were seeded and infected as described previously. For calculation of percentage of THP1 cells infected, in/out staining was performed as follows. 40 minutes post addition of bacteria, T=0, cells were washed three times with cold PBS. Rabbit anti-*sonnei* (1:100) (phase 1 and 2 sera, Fisher Scientific) or rabbit anti-flexneri (1:500) (serotype 5a sera, PHE) diluted in 2% Bovine serum albumin (BSA):PBS were added to cells. Cells were incubated with antibodies on ice for 30 minutes. Following this, cells were washed with cold PBS, and incubated on ice with donkey anti-rabbit-Alexa594. (1:500, 2% BSA:PBS). Cells were fixed with 2% paraformaldehyde (PFA) diluted in PBS for 20 minutes, washed in PBS, and neutralised with 50 mM NH_4_Cl. 0.1% Triton-X 100 was added to cells for 8 minutes to permabilise. DAPI (1:1000, 2% BSA:PBS) (Invitrogen) and phalloidin-Alexa647 (1:100) (Invitrogen) were incubated with cells. Coverslips were mounted onto slides with ProLong® Gold Antifade Mountant and visualised using Zeiss Axio Observer Z1 microscope. For counting of ASC foci, ASC^mRFP^ cells (46) were infected as described, washed in PBS at 3 hours post infection and fixed. The protocol was then continued as described above.

### LPS preparation and visualisation

Crude LPS was prepared as follows. 1.5 ml of overnight culture was centrifuged, resuspended in laemmli buffer and boiled for 5 min. Proteinase K (1 mg/ml) was added and incubated for 2 h at 56°C. 2-mercaptoethanol (5%) was added and samples were boiled for 5 min and 5 ul of each sample separated by 12% SDS-PAGE. The gel was either transferred to PVDF and incubated with *S. flexneri* serotype 5a antibody (PHE) or *S. sonnei* phase I antibody (Abcam), followed by anti-rabbit HRP and developed by chemiluminescence, or fixed and silver stained as previously described (53).

### Statistical Analysis

The number of independent repeats performed for each experiment is indicated (by n) in the figure legend. One-way ANOVA or Student’s t-test was performed to compare means as implemented in GraphPad prism 8. Errors bars represent standard error of the mean throughout.

## Acknowledgements

ARS would like to acknowledge funding from the MRC (MR/P022138/1). JLW is the recipient of an MRC Centre for Molecular Bacteriology and Infection (CMBI) PhD scholarship as part of the CMBI Centre Award MR/J006874/1.

---

**Figure S1.**
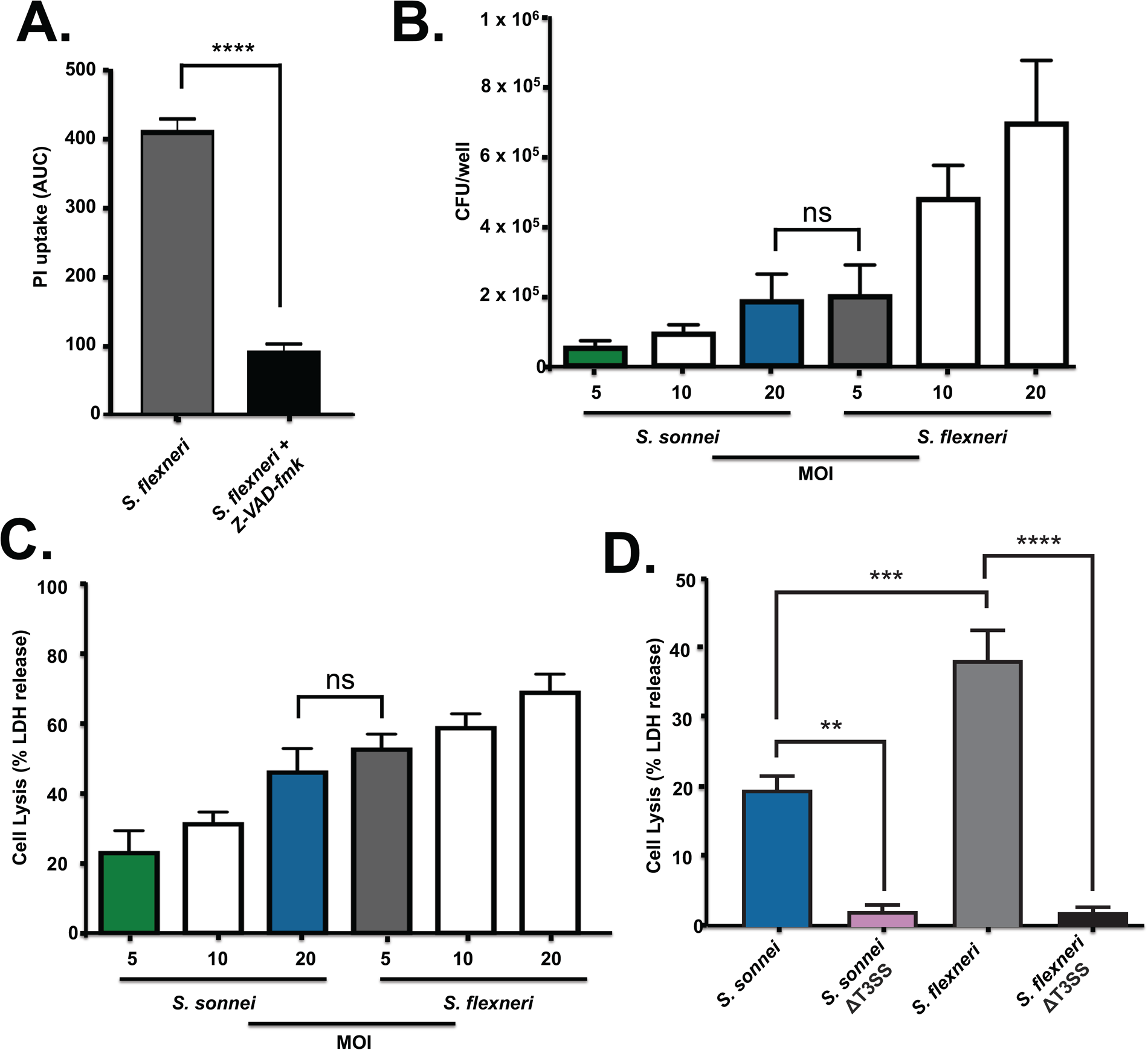
(A) THP1 cells were left untreated or treated with 50 μM Z-VAD-fmk one hour prior to infection with *S. flexneri*. Cell death was measured by PI uptake over a 3 h time-course and plotted as Area Under the Curve (AUC). ****p<0.0001 by unpaired Student’s t test, n=3 (B) THP1 cells were infected with the indicated MOIs of *S. sonnei* or *S. flexneri* and the number of cytosolic CFU were enumerated after treating cells with gentamicin and chloroquine to kill extracellular and vacuolar bacteria. Ns, non-significant by one-way ANOVA, n=4. (C) THP1 cells were infected with wild type *S. sonnei* and *S. flexneri* and their respective T3SS mutants. Cell death was measured by LDH release at 3 h. **p<0.01, ***p<0.001, ****p<0.0001 by one-way ANOVA, n=3. (D) THP1 cells were infected with *S. sonnei* or *S. flexneri* and the respective T3SS mutants. Cell death was measured by LDH release at 3 h. ns, non-significant by one-way ANOVA, n=4.

**Figure S2.**
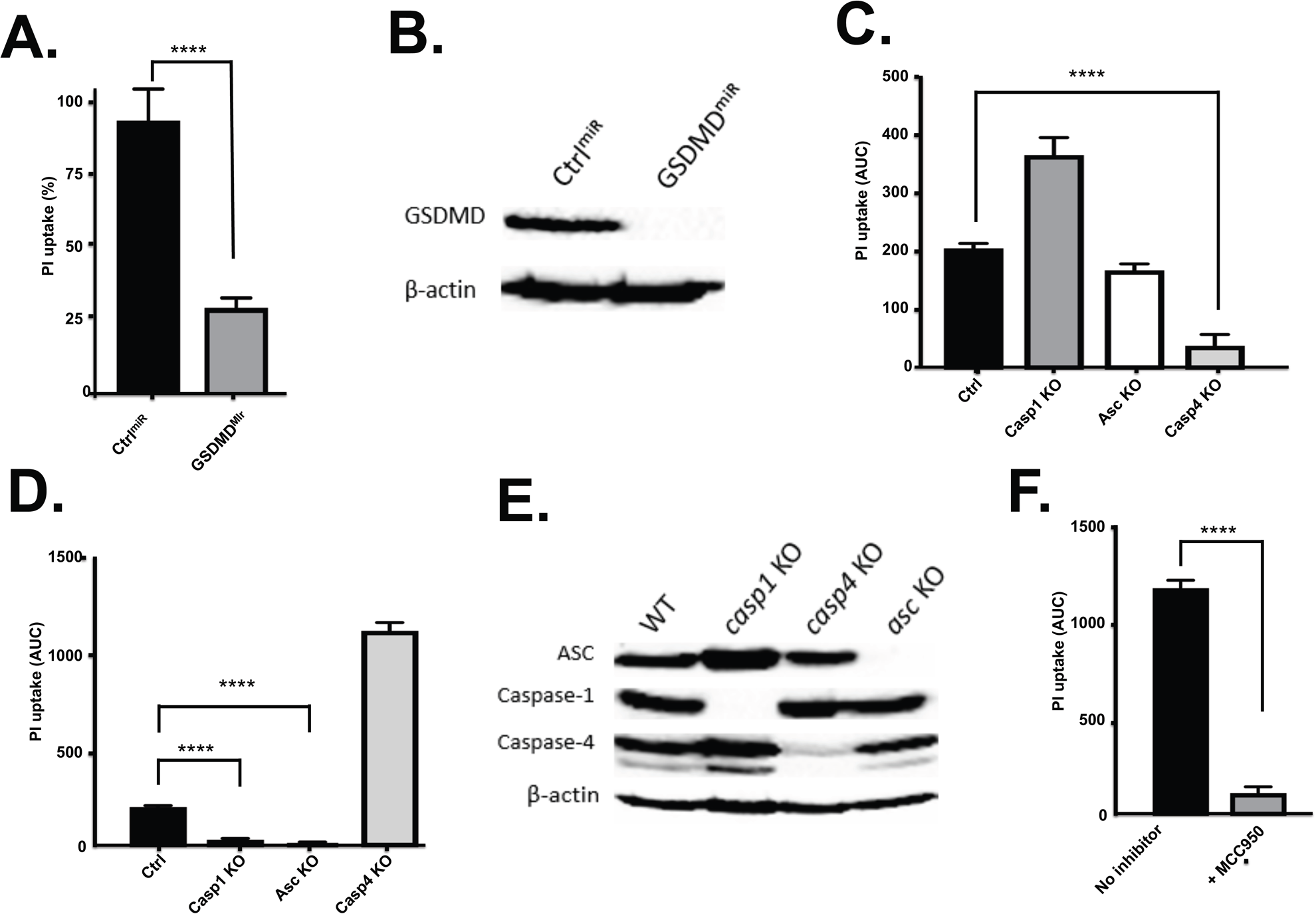
Phenotypic confirmation of THP1 cell lines. (A) GSDMD^miR^ and Control^miR^ THP1 cells were treated with LPS and nigericin. Cell death was measured by PI uptake over a 3 h time-course and plotted as Area Under the Curve (AUC). ****p<0.0001 by one-way ANOVA, n=3. (B) Representative immunoblots show silencing of GSDMD, β-actin serves as a loading control. (C and D) Control THP1 cells and THP1 cells deficient for caspase-1, ASC or caspase-4 were transfected with LPS to induce caspase-4 activation (D) or LPS + nigericin to induce canonical NLRP3 activation (E). Cell death was measured by PI uptake over a 3 h timecourse and plotted as AUC. ****p<0.0001 by one-way ANOVA, n=3. (E) Representative immunoblot shows knockout of Caspase-1, ASC or Caspase-4 loss in the appropriate cell line. β-actin serves as a loading control. (F) LPS + nigericin was added to THP1 cells incubated with or without 5 μM MCC950. Cell death was measured by PI uptake over a 3 h timecourse and plotted as AUC. ****p<0.0001 by one-way ANOVA, n=4.

**Figure S3.**
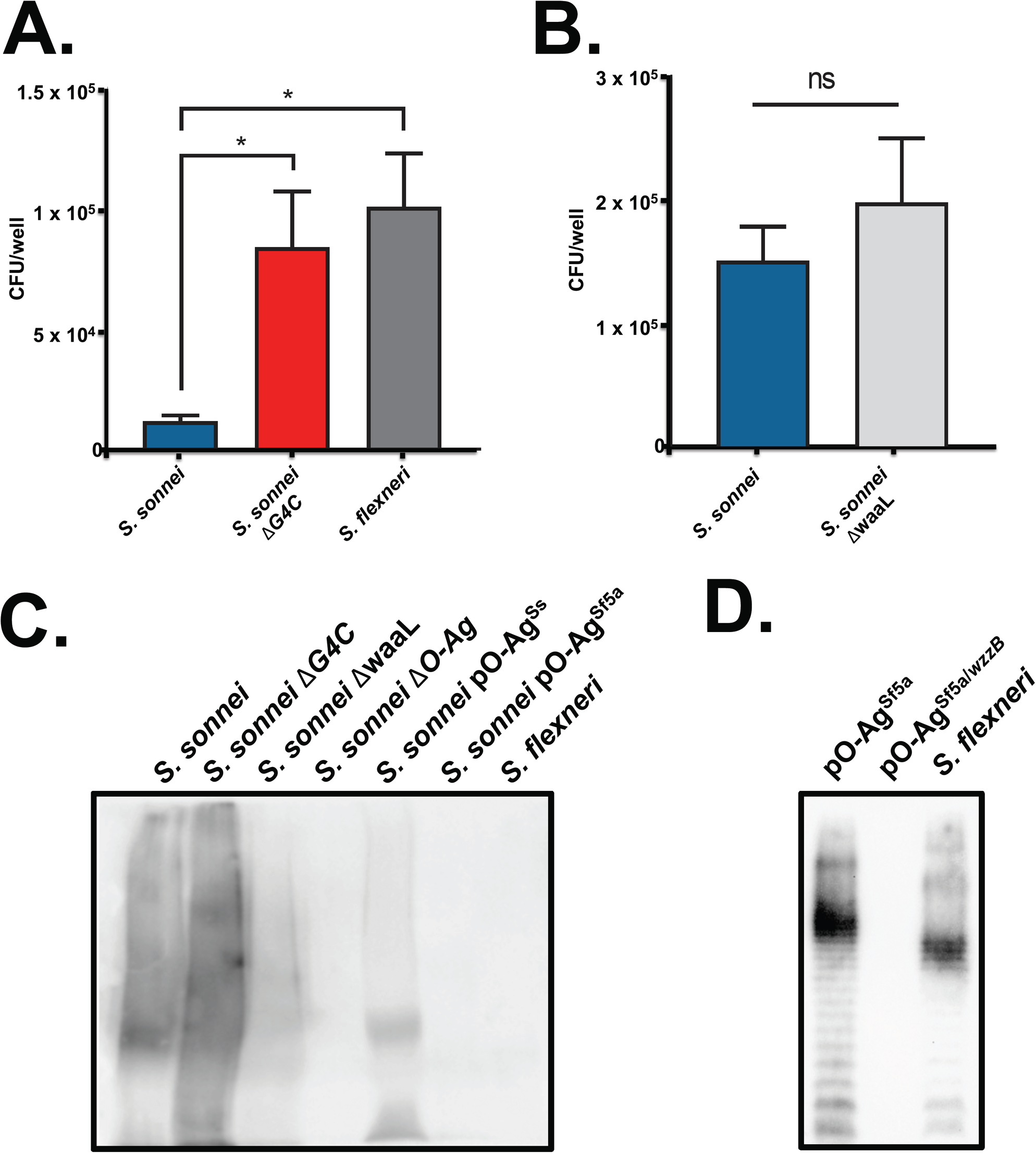
Characterisation of *S. sonnei* O-Ag mutants. (A) HeLa cells were infected with *S. sonnei, S. sonnei* Δg4c or *S. flexneri* and gentamicin protected internalised bacteria were determined by CFU enumeration. *p<0.05 by one-way ANOVA, n=3. (B) THP1 cells were infected with wild type *S. sonnei* or *S. sonnei* Δ*waaL* and gentamicin-protected internalised bacteria were determined by CFU enumeration. Ns, non-significant by one-way ANOVA, n=5 (C and D) Crude LPS was purified from the indicated *S. sonnei* and *S. flexneri* strains and the presence of the *S. sonnei* O-Ag (C) or *S. flexneri* 5a O-Ag (D) was detected using serotype specific antibodies.

**Table S1:**
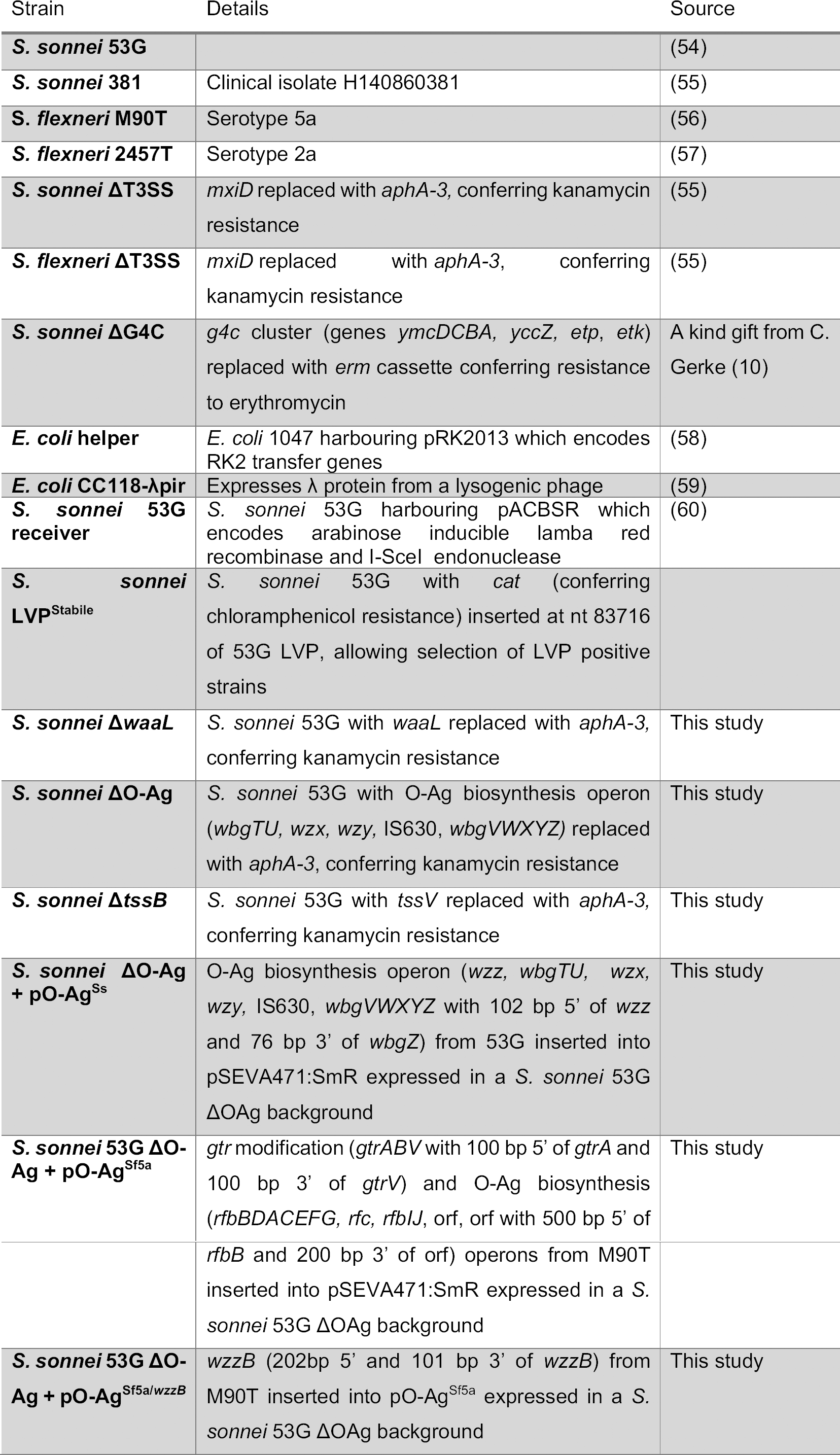
Strains used in this study

**Table S2:**
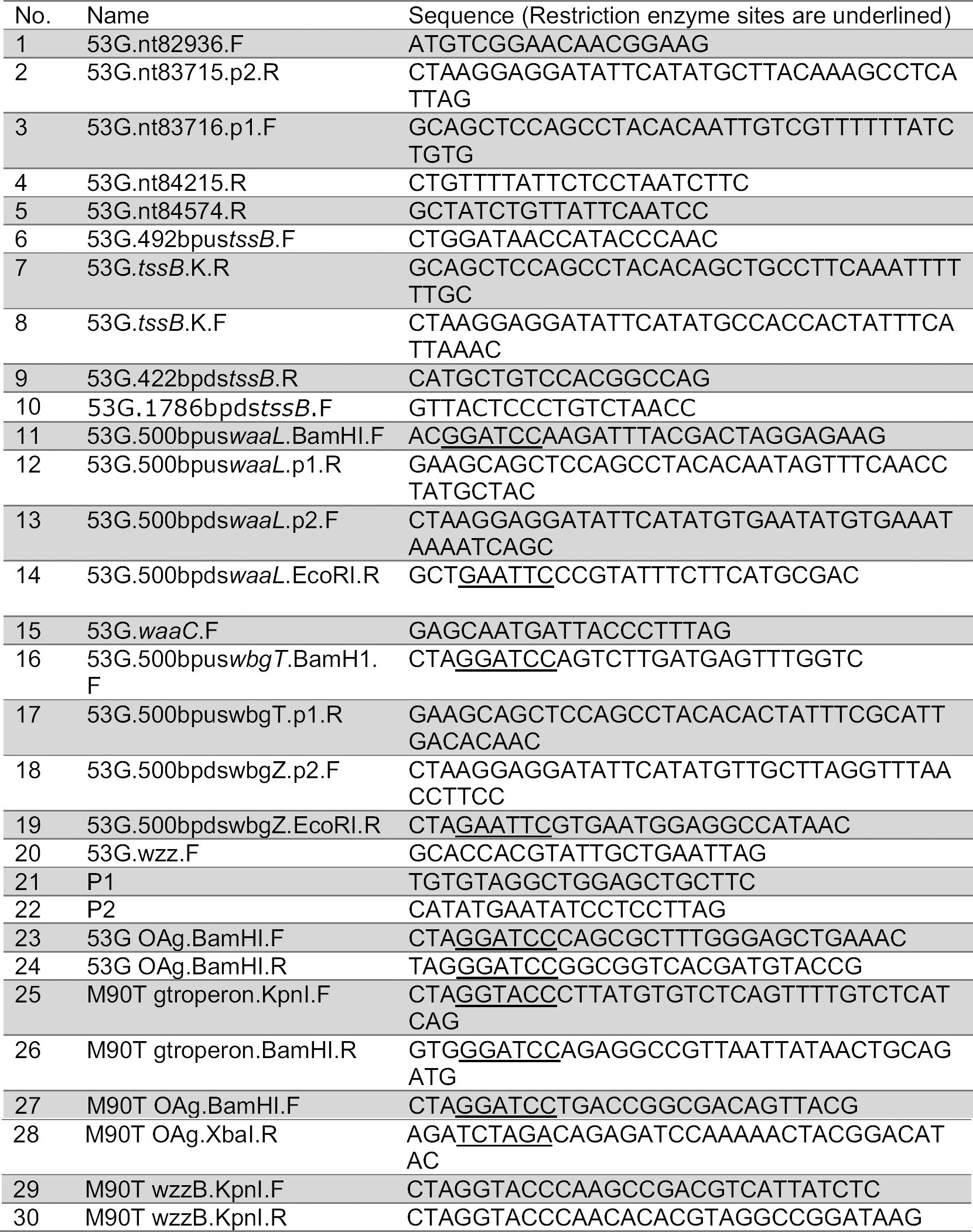
Primers used in this study.

